# Cortisol Reactivity and Secondary Hyperalgesia induced by High-Frequency Electrical Stimulation: The Influence of Romantic Partner Touch

**DOI:** 10.64898/2026.11.26.769416

**Authors:** Márcia da-Silva, Anabela Silva-Fernandes, Adriana Sampaio, Joana Coutinho, Alberto J. González-Villar

**Affiliations:** Psychological Neuroscience Laboratory (PNL), Research Center in Psychology (CIPsi), School of Psychology, University of Minho, Braga, Portugal; Center for Psychology at University of Porto, Faculty of Psychology and Education Sciences – University of Porto, Rua Alfredo Allen, 4200–135 Porto, Portugal; Instituto de Psicoloxía (IPsiUS), Universidade de Santiago de Compostela, Santiago de Compostela, Spain

**Keywords:** Salivary cortisol, Secondary hyperalgesia, Affective touch, Social context, HPA axis

## Abstract

Romantic partner touch has been shown to have analgesic properties, yet its influence on neuroendocrine mechanisms associated with pain sensitization remains unclear.

This study investigated whether romantic partner’s touch modulates salivary cortisol responses during an experimental pain protocol and whether baseline cortisol levels or cortisol reactivity are associated with experimentally induced secondary hyperalgesia. Sixty-four adults completed two counterbalanced sessions - one alone and one accompanied by their romantic partner. In each session, participants provided a baseline saliva sample, underwent high-frequency electrical stimulation to induce secondary hyperalgesia, and provided a second saliva sample at the end of the protocol. In the accompanied session, participants received their partner’s affective touch during and after stimulation.

Baseline cortisol levels did not differ between sessions and were not associated with secondary hyperalgesia. Cortisol concentrations decreased from baseline to post-task, with Bayesian analysis supporting this decrease. The session × sampling time interaction was not significant. Cortisol reactivity was not associated with secondary hyperalgesia in either condition. Furthermore, between-session differences in cortisol reactivity were not associated with between-session differences in secondary hyperalgesia. Bayesian analyses favored the null models for these associations.

These findings provide no evidence that romantic partner touch modulated changes in salivary cortisol concentrations during the experimental pain protocol. Moreover, salivary cortisol assessed before and after the protocol was not associated with experimentally induced secondary hyperalgesia, suggesting that these measures may not capture HPA-axis processes associated with the development of pain sensitization under the present experimental conditions.

## 1. Introduction

Affective touch has received growing attention as a contextual modulator of pain and stress. Gentle stroking at CT-optimal velocities preferentially activates C-tactile afferents (CT), which are associated with the affective dimension of touch and with pleasant somatic experience (Ackerley et al., 2014; Löken et al., 2009; Marshall & McGlone, 2020; McGlone et al., 2014). Beyond its sensory properties, affective touch occurring within socially meaningful contexts, for example, between romantic partners, and in the presence of social support, has been associated with reduced subjective and electrophysiological responses to pain (da-Silva et al., 2026a; Krahé et al., 2015; von Mohr et al., 2018). Although the underlying biological mechanisms remain unclear, one possible explanation is that these effects are partly mediated by stress regulation processes. Cortisol, a hormone commonly used as a marker of hypothalamic-pituitary-adrenal (HPA) activity in response to stress in humans, offers a useful way to examine this possibility. Cortisol, also known as the “stress hormone,” is a glucocorticoid produced in the zona fasciculata of the adrenal cortex following activation of the hypothalamic– pituitary–adrenal (HPA) axis, with a steroid structure derived from cholesterol. Because salivary cortisol provides a non-invasive measure of physiological stress responses and can be repeatedly sampled throughout experimental protocols, it has become one of the most widely used biomarkers in human psychophysiological research.

Social support and forms of physical contact, such as embracing, have been shown to attenuate cortisol responses to psychosocial stress (Berretz et al., 2022; Dreisoerner et al., 2021; Heinrichs et al., 2003; Hostinar & Gunnar, 2015). Evidence specifically examining CT-targeted affective touch as modulating stress response is more limited. In infants, repeated CT-optimal stroking has been associated with reduced physiological stress responses, including differences in cortisol regulation (Van Puyvelde et al., 2021), while a prolonged CT-targeted affective touch intervention in institutionalized and fostered children was associated with normalization of salivary cortisol levels (Nikolaeva et al., 2024). In adults, however, Triscoli et al., (2017).found no specific effect of CT-optimal stroking on salivary cortisol compared with vibrotactile stimulation.

In pain research, salivary cortisol has been widely used to characterize neuroendocrine responses to nociceptive stimulation. However, findings have been inconsistent, with changes being reported more consistently following cold pain than after heat, mechanical, or visceral pain paradigms (Goodin et al., 2012; Zarnegar et al., 2024, 2025). Furthermore, salivary cortisol responses depend on the intensity of the stimulus and have been shown to peak approximately 10–20 minutes after pain induction (Goodin et al., 2012; Zarnegar et al., 2025). Given that salivary cortisol responses peak with some delay following painful stimulation, post-testing cortisol levels may be better suited to reflect the overall neuroendocrine response to the pain-testing session (the cumulative physiological impact of a pain protocol) than the isolated, brief stimulus-specific effects (Quartana et al., 2010).

Beyond characterizing cortisol responses to pain, experimental studies suggest that glucocorticoid activity may also influence pain processing, although the direction and magnitude of these effects appear to depend on the pain modality, sex, and experimental context. Pharmacological interventions that alter cortisol levels or involve glucocorticoid administration have been reported to increase, attenuate, or leave unchanged experimental pain sensitivity (Benson et al., 2019; Kuehl et al., 2010; Michaux et al., 2012; Wingenfeld et al., 2015). These findings indicate that HPA-axis activity may modulate pain processing, but that its effects may depend on the characteristics of the pain model.

Of particular relevance to the present study is whether HPA-axis activity is also related to pain sensitization. Experimental manipulations of cortisol have produced inconsistent effects on temporal summation of pain (Bruehl et al., 2022; Kuehl et al., 2010; Michaux et al., 2012). Secondary hyperalgesia (SH), commonly quantified as the area of mechanical hypersensitivity surrounding the site of injury, is considered a well-established marker of central sensitization in humans and provides a useful experimental model for addressing this question (Leone et al., 2025; Woolf, 2011). Glucocorticoid activity, life stress and social support have all been reported to modulate SH, although not always its spatial extent (Jaltare et al., 2023, 2024; Michaux et al., 2012; You et al., 2016). It nevertheless remains unclear whether baseline cortisol levels or the overall cortisol response to an experimental pain protocol are associated with the magnitude of secondary hyperalgesia, and whether these associations differ according to the social context in which pain is experienced. Therefore, the present study aimed to investigate whether baseline cortisol levels and changes in cortisol protocol response are related to the magnitude of experimentally induced secondary hyperalgesia and whether the presence and touch of a romantic partner modulate salivary cortisol responses during an experimental pain protocol.

We hypothesized that the presence and touch of a romantic partner would influence both baseline cortisol levels and cortisol reactivity during the experimental pain protocol. We further hypothesized that both baseline cortisol levels and cortisol reactivity would be associated with the magnitude of experimentally induced secondary hyperalgesia.

## 2. Methods

### 2.1. Ethics

The study was conducted in accordance with the Declaration of Helsinki and approved by the Ethics Committee for Social and Human Sciences of the University of Minho (CEICSH 030/2022). All participants provided informed consent prior to participation, confirming that their involvement was voluntary and that they could withdraw from the study at any time.

### 2.2. Participants

Sixty-four participants (thirty-two couples; aged 18-41 years, M = 25.3, SD = 5.7; 33 males, 31 females) were originally enrolled in the study. For the present study, cortisol data were available for 57 participants (28 couples: aged 18-37 years, M=24.8, SD=5.3; 29 males, 28 females), who were included in the analyses. Couples were recruited via the University of Minho Credits platform, social media, advertising flyers, and a snowball sampling approach. Eligibility criteria required both partners to be in a stable romantic relationship for at least one year. Exclusion criteria included a history of cardiac, psychiatric, or neurological disorders, chronic pain, substance abuse, pain on the day of testing, or any skin injuries on the forearms or hands. Participants were not informed about the study’s hypotheses. University students received course credits, while non-student participants were compensated with vouchers worth €60.

### 2.3. Procedure

Upon arrival at the laboratory, participants were briefed on the study procedures and provided written informed consent.

The study comprised two counterbalanced sessions-an accompanied session and an alone session-conducted at least one week apart to minimize carry-over effects (see figure 1). In each session, participants were comfortably seated in a soundproofed, light-controlled room and completed a 4-minute resting-state period recorded with eyes open. In the alone session, no other person was present in the room. In the accompanied session, the participant’s romantic partner was seated next to the participant, but no physical contact or verbal interaction was permitted. The experimenter remained in an adjacent room and entered only to set up the equipment, explain the task, and perform the pinprick mapping, following the same procedure in both sessions.

**Figure 1.**
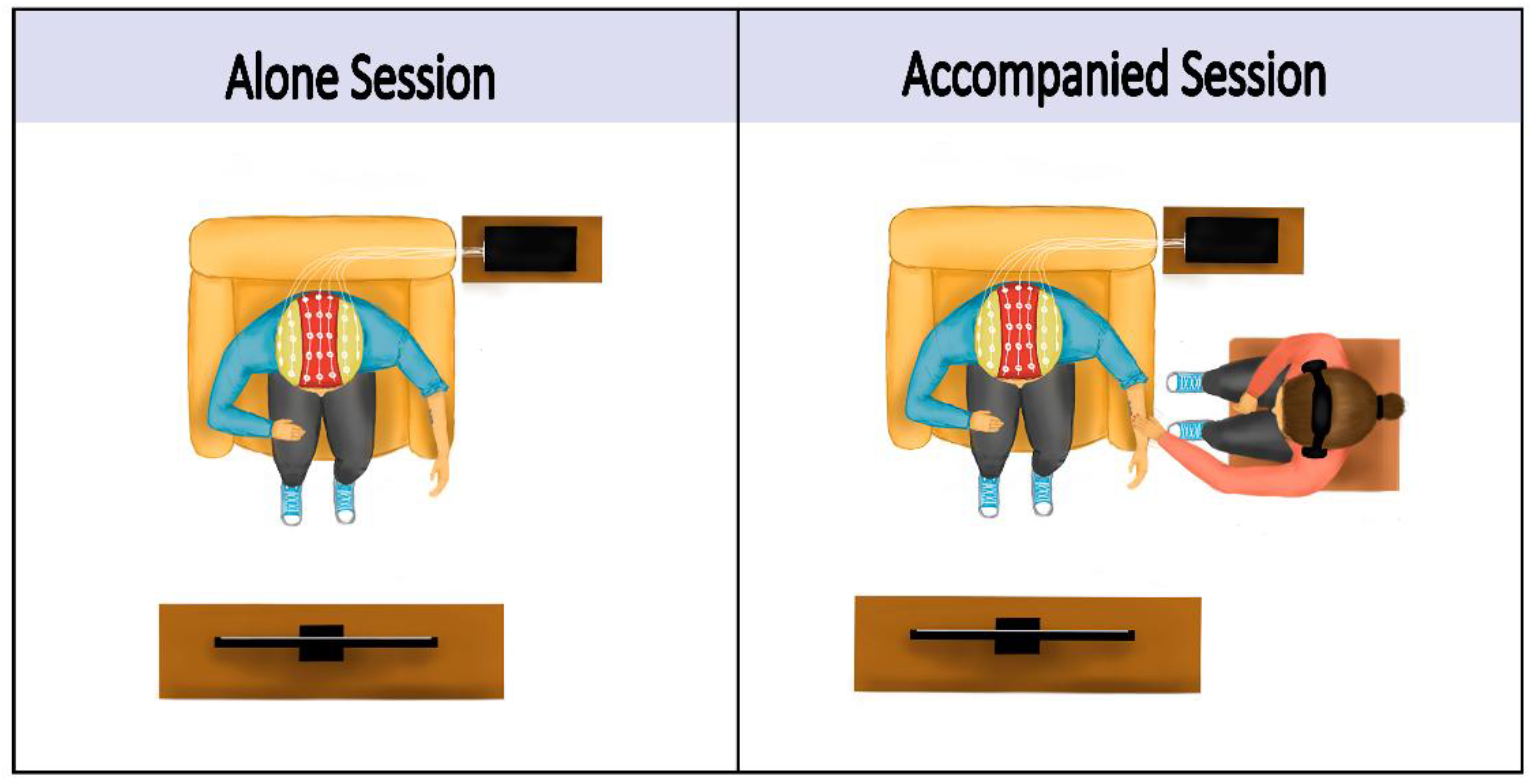
Experimental conditions during the alone and accompanied sessions. The figure illustrates the two experimental conditions used during the SH protocol. In both sessions, participants underwent high-frequency stimulation (HFS) followed by a 10-minute post-stimulation period. In the accompanied session, the participant’s romantic partner was present in the room and delivered affective touch during HFS and during the 10 min post-HFS, whereas in the alone session, no partner was present and no touch was administered.

Immediately following the resting-state recording, participants provided a saliva sample for cortisol analysis (baseline cortisol levels). Participants then completed an additional experimental task, investigating the effects of affective CT-touch on temporal summation of second pain (TSSP), which has been reported separately (da-Silva et al., 2026a) and is unrelated to the present analyses, before undergoing the Secondary Hyperalgesia (SH) protocol. The SH protocol, also described in (da-Silva et al., 2026b) was induced using high-frequency electrical stimulation (HFS) consisting of five 1-second trains of electrical pulses at 100 Hz applied to the left volar forearm, at an intensity of 20X the detection threshold. After HFS administration, participants remained at rest for 10 minutes. In the accompanied session, during HFS, affective touch was applied to the left dorsal forearm, in alternating 5-second periods of stroking and 5-second periods of no-touch. The first stroking bout began 1 second after the onset of the first train of electrical pulses, and the last ended 6 seconds after the final train. During the subsequent 10-minute post-HFS period, affective touch was delivered for 30 seconds, followed by 20-second no-touch intervals. Auditory cues were presented every 3.5 seconds to maintain a stroking velocity of approximately 4 cm/s, which falls within the CT-optimal range of approximately 3–5 cm/s (Ackerley et al., 2014; Löken et al., 2009). During the alone session, participants remained alone during the HFS administration and for 10 minutes afterwards.

Secondary hyperalgesia was quantified as the total area of mechanical hypersensitivity (cm²) derived from pinprick mapping using a calibrated 128 mN probe following high-frequency stimulation. A saliva sample was collected at the end of the SH protocol, approximately 17–20 minutes after HFS.

### 2.4. Measures

#### 2.4.1. Enzyme-linked immunosorbent assay of cortisol levels

Saliva samples were collected using Salivette devices (Sarstedt, Germany) and centrifuged for to recover the saliva for 10 min at 13000 rpm using a Gyrozen 1730R microcentrifuge at 4°C to remove debris. The samples were then stored at −80 °C until analysis. Cortisol levels were quantified, all in the same day, following the manufacturer’s instructions (Demeditec Diagnostics, Germany, cat. no. DES6611). All samples, calibrators, and controls were assayed in duplicate, with 50 μL dispensed into the appropriate wells.

Optical density was measured at 450 nm within 15 minutes using an ELISA plate reader from Bio-Rad (model iMark). Data preparation was performed in Microsoft Excel by calculating the mean optical density for each set of calibrators, controls, and samples. A standard curve was generated by plotting optical density values against the logarithm of cortisol concentrations. Sample concentrations were determined by interpolation from the standard curve.

Intra-assay coefficient of variability was calculated to be 4.1%; inter-assay coefficient of variability was calculated to be 3.3% for low-control and 4.5% for high-control.

#### 2.4.2- Area of Secondary hyperalgesia

Pinprick stimulation was applied at approximately 1-cm intervals along four linear trajectories extending from the wrist and cubital fossa (proximal–distal axis) and from the lateral and medial edges of the forearm (lateral–medial axis) toward the center of the HFS stimulation site. Stimulation along each trajectory continued until participants reported a noticeable change in pinprick intensity, indicating the boundary of the hyperalgesic area. These four boundary points were marked on the skin with a makeup pencil and the corresponding distances were measured using a digital caliper. The area of SH was then estimated by calculating the area of the diamond defined by the four boundary points.

### 2.5. Statistical Analysis

All statistical analyses were conducted in JASP (version 0.19.0; JASP Team, 2024). Model assumptions were assessed through visual inspection of residual and Q–Q plots, and boxplots were examined to identify potential outliers. One participant was excluded from all analyses due to being identified as an outlier.

To examine the effect of social context on baseline cortisol levels, a repeated-measures ANOVA was conducted with session (accompanied vs alone) as a within-subject factor, sex as a between-subject factor, and baseline cortisol levels as the dependent variable.

To control for potential confounding effects of circadian variation in cortisol secretion, a binary covariate was created to indicate whether cortisol samples were collected at comparable times across sessions (0 = same time of day across sessions; 1 = different times of day between sessions). This variable was included as a covariate in analyses of absolute cortisol concentrations to account for potential diurnal influences on cortisol levels. It was not included in analyses of cortisol reactivity, as reactivity was calculated as the within-session change from baseline to post-task cortisol levels.

To examine changes in cortisol concentrations between baseline and post-task levels, a 2×2 repeated-measures ANOVA was conducted with session (accompanied vs alone) and sampling time (baseline vs post-task) as repeated measures factors, and sex as a between-subjects factor. The time-of-day control variable was included as a covariate to account for potential differences in the timing of cortisol sampling between sessions.

Cortisol reactivity was quantified as the change in cortisol concentration across the experimental protocol (Δ cortisol = post-protocol concentration − baseline concentration). Differences in cortisol reactivity between accompanied and alone sessions were examined using repeated-measures ANOVA with session (accompanied vs. alone) as the within-subject factor and sex as the between-subject factor.

To investigate the relationship between cortisol and experimentally induced secondary hyperalgesia, separate multiple linear regression analyses were performed for the accompanied and alone sessions. Secondary hyperalgesia area was entered as the dependent variable. Baseline cortisol levels or cortisol change (Δ cortisol), together with relationship duration, were entered as independent variables in separate regression models. Statistical significance was set at *p* < .05.

Lastly, Bayesian analyses were conducted to complement the frequentist analyses and assess the relative evidence for the alternative versus null hypotheses. Bayes factors (BF₁₀) were computed, with BF₁₀ values quantifying the evidence in favor of the alternative hypothesis relative to the null hypothesis. Bayesian repeated-measures ANOVAs were conducted in JASP, using Cauchy priors for effect sizes (scale = 0.5 for fixed effects and 1.0 for random effects). Bayesian linear regression analyses were also conducted in JASP using the JZS prior (r scale = 0.354) and a uniform model prior (Beta-binomial distribution, a = 1, b = 1).

## 3. Results

### 3.1 Baseline Cortisol levels

Descriptive statistics for baseline and post-task cortisol levels are presented in Table 1. We did not observe a significant main effect of session on baseline cortisol levels (F(1, 53) = 0.956, *p* = .33, h_p_^2^= 0.018, BF_10_=0.74), indicating that cortisol levels did not differ between accompanied and alone conditions at the baseline levels. There was also no significant interaction between session and sex (F(1, 53) = 2.859, *p* = .097, h_p_^2^= 0.051, BF_10_=0.64), nor between session and the time-of-day covariate (F(1, 53) = 0.381, *p* = .540, h_p_^2^= 0.007, BF_10_=0.36).

**Table 1.**
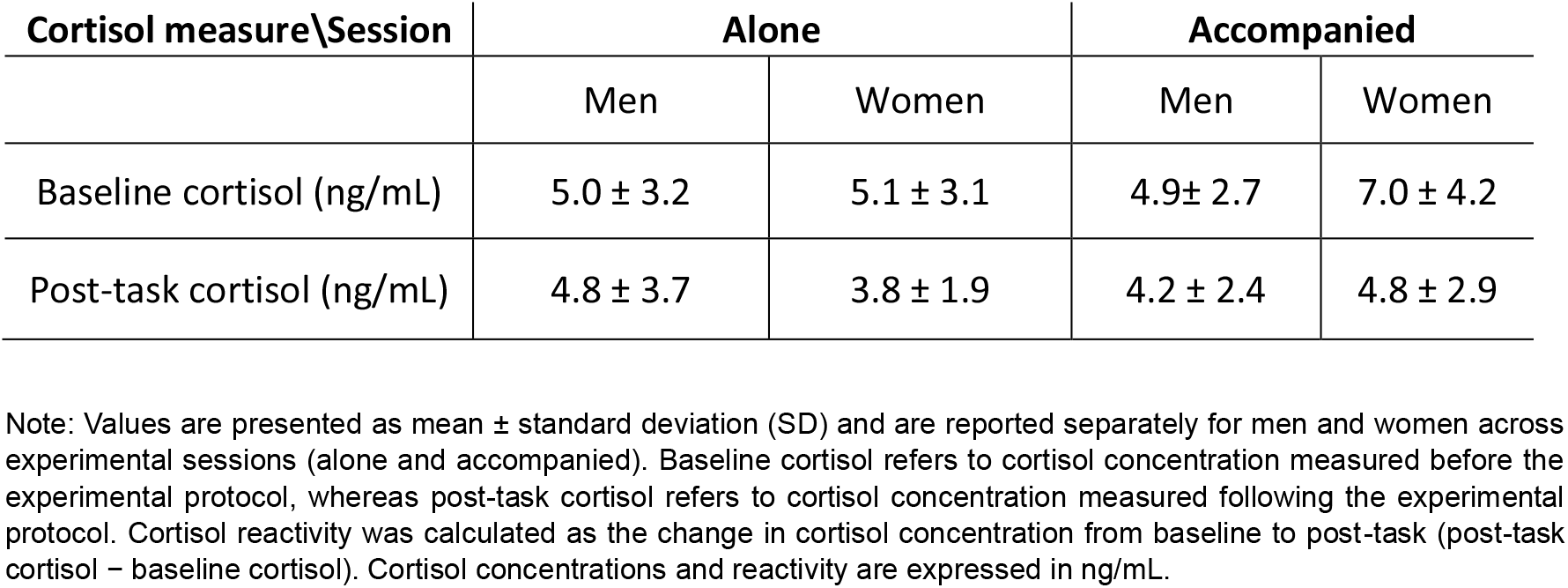
Descriptive statistics of cortisol concentrations and cortisol reactivity across experimental sessions.

Regarding between-subject effects, the main effect of sex was not statistically significant (F(1, 53) = 2.99, *p* = .090, h_p_^2^= 0.053, BF_10_=0.86). Similarly, the covariate controlling for time-of-day differences was not statistically significant (F(1, 53) = 0.146, *p* = .707, h_p_^2^= 0.003, BF_10_=0.47).

### 3.2 Cortisol levels across sampling times

A two-way repeated-measures ANOVA examining the effects of session (alone vs. accompanied) and sampling time (baseline vs. post-task), with sex included as a between-subjects factor, revealed a significant main effect of sampling time (*F*(1, 54) = 10.548, *p* = .002, ηp² =.163, BF_10_=9.33), indicating that cortisol concentrations decreased from baseline to post-task across sessions and sexes (see figure 2 and table 1). There was no significant main effect of session, (*F*(1, 54) = 1.697, *p* = .198, ηp² = .030, BF_10_=0.47) , indicating that cortisol concentrations did not differ significantly between the accompanied and alone conditions. The main effect of sex was also not significant, (*F*(1, 54) = 0.759, *p* = .387, ηp² = .014, BF_10_=0.35). However, there was a significant session × sex interaction (*F*(1, 54) = 4.274, *p* = .043, ηp² = .073), suggesting that the effect of session on cortisol concentrations differed by sex. In contrast, the time × sex interaction did not reach statistical significance, (*F*(1, 54) = 3.841, *p* = .055, ηp² = .066), and neither the session × time interaction, (*F*(1, 54) = 1.393, *p* = .243, ηp² = .025) nor the three-way session × time × sex interaction, (*F*(1, 54) = 0.169, *p* = .683, ηp² = .003), was significant. Bayesian model including sampling time, sex, and their interaction yielded *BF*₁₀ = 4.020, while the model including session, sampling time, sex, and the session × sex and time × sex interactions yielded *BF*₁₀ = 2.862. Importantly, time of day did not significantly interact with session (F(1, 53) = 1.646, p = .205, ηp² = .030) or with sampling time (F(1, 53) = 5.319 × 10⁻⁴, p = .982, ηp² < .001) indicating that the effects of session and sampling time were not significantly influenced by the time of day at which cortisol was assessed.

**Figure 2.**
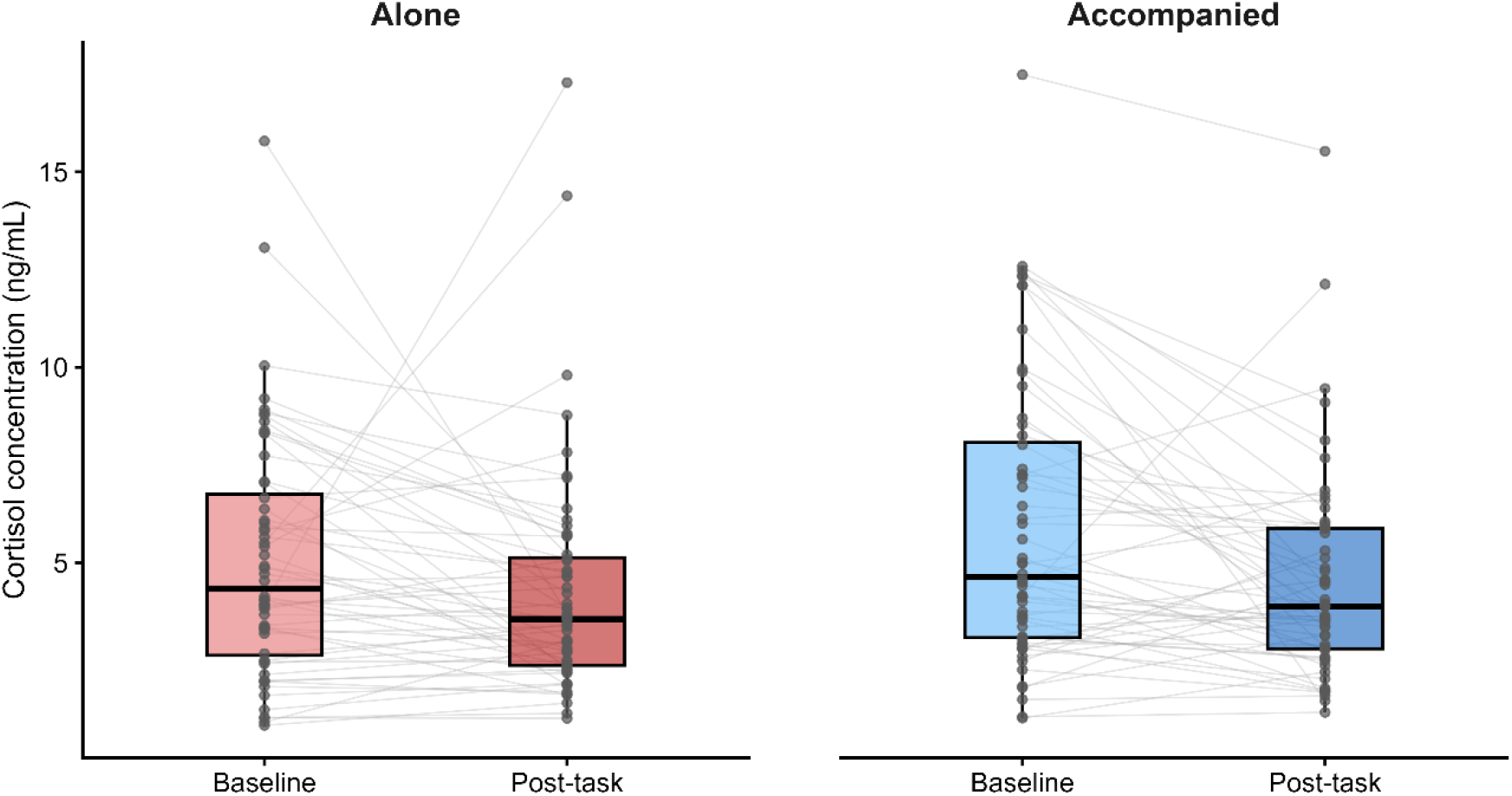
Cortisol concentrations at baseline and post-task during the alone and accompanied sessions. The left panel shows cortisol concentrations during the alone session, and the right panel shows cortisol concentrations during the accompanied session. Boxplots represent the median and interquartile range, grey points represent individual participant values, and paired observations from the same participant are connected by grey lines.

### 3.3 Baseline cortisol and secondary hyperalgesia

#### 3.3.1 Alone Session

A multiple linear regression was performed to examine whether baseline cortisol levels and relationship duration predicted secondary hyperalgesia area during the alone session (see figure 3A). The overall model was not significant, F(2, 53) = 0.053, p = .948, explaining 0.2% of the variance in secondary hyperalgesia area (R² = .002, adjusted R² = −.036), BF₁₀ = 0.11. Baseline cortisol levels were not a significant predictor of secondary hyperalgesia area (B = −0.090, SE = 0.569, β = −0.022, t(53) = −0.158, p = .875, BF₁₀ = 0.28). Relationship duration was also not a significant predictor (B = −0.010, SE = 0.035, β = −0.040, t(53) = −0.292, p = .771, BF₁₀ = 0.27).

**Figure 3.**
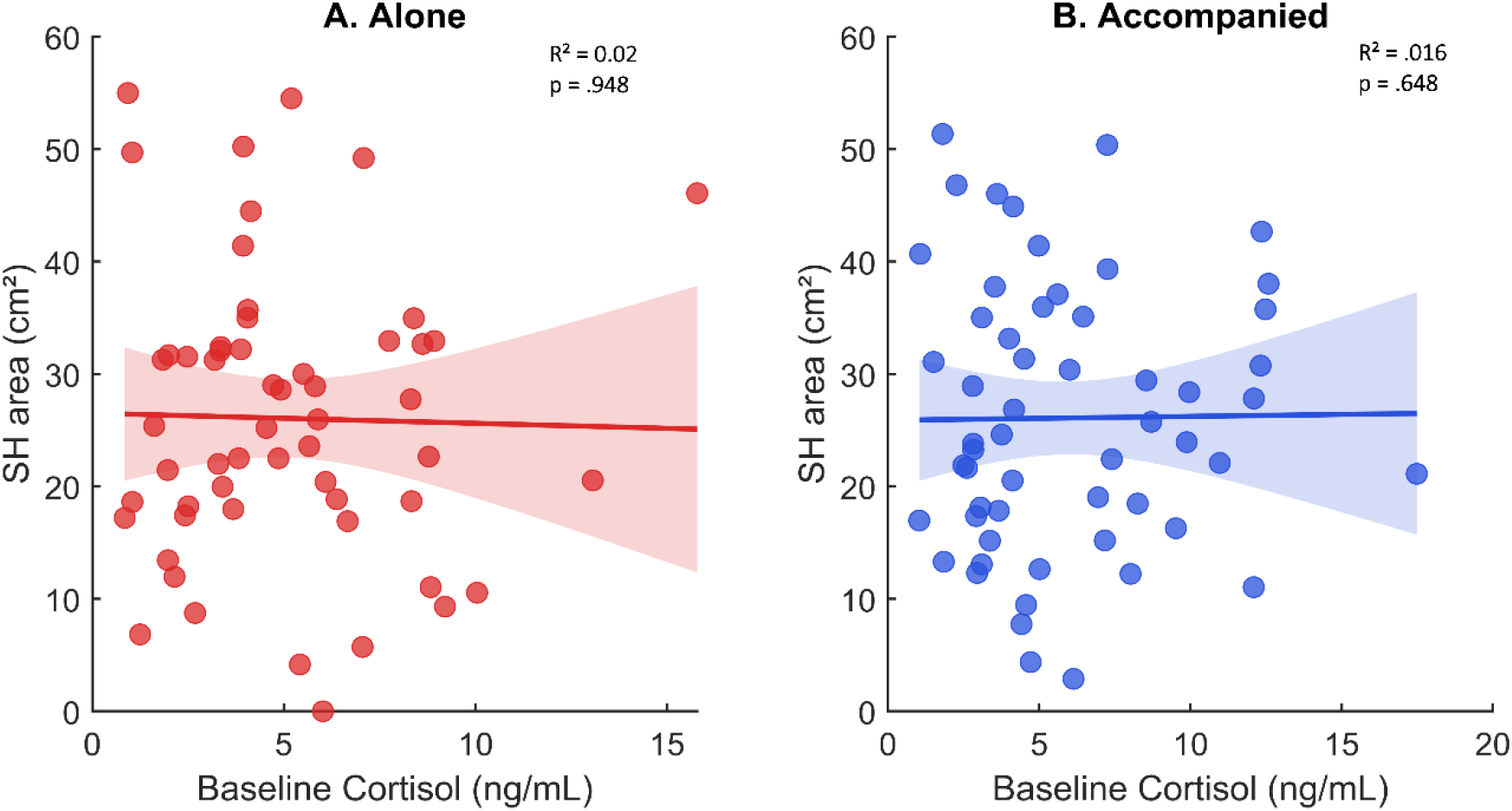
Scatterplots show the association between baseline cortisol levels and secondary hyperalgesia area during the alone (A) and accompanied (B) sessions. Each point represents one participant. Regression lines are adjusted for relationship duration, with shaded areas indicating 95% confidence intervals. R² and *p*-values shown in the figure correspond to the overall multiple regression models including baseline cortisol and relationship duration as predictors of secondary hyperalgesia area.

#### 3.3.2 Accompanied Session

A multiple linear regression was performed to examine whether baseline cortisol levels and relationship duration predicted secondary hyperalgesia area during the accompanied session (see figure 3B). The overall model was not significant, F(2, 53) = 0.438, p = .648, explaining 1.6% of the variance in secondary hyperalgesia area (R² = .016, adjusted R² = −.021), BF₁₀ = 0.15. Baseline cortisol levels were not a significant predictor of secondary hyperalgesia area (B = 0.035, SE = 0.443, β = 0.011, t(53) = 0.079, p = .937, BF₁₀ = 0.27). Relationship duration was also not a significant predictor (B = 0.030, SE = 0.032, β = 0.127, t(53) = 0.933, p = .355, BF₁₀ = 0.39).

### 3.4 Cortisol reactivity and secondary hyperalgesia

#### 3.4.1 Alone Session

A multiple linear regression was performed to examine whether cortisol reactivity and relationship duration predicted secondary hyperalgesia area during the alone session (see figure 4A). The overall model was not significant, *F*(2, 53) = 0.143, *p* = .867, explaining 0.5% of the variance in secondary hyperalgesia area (*R*² = .005, adjusted *R*² = −.032) BF₁₀ = 0.12. Cortisol reactivity was not a significant predictor of secondary hyperalgesia area (*B* = 0.228, *SE* = 0.504, β = 0.062, *t*(53) = 0.451, *p* = .654, BF₁₀ = 0.30). Relationship duration was also not a significant predictor (*B* = −0.010, *SE* = 0.035, β = −0.038, *t*(53) = −0.279, *p* = .781, BF₁₀ = 0.27).

**Figure 4.**
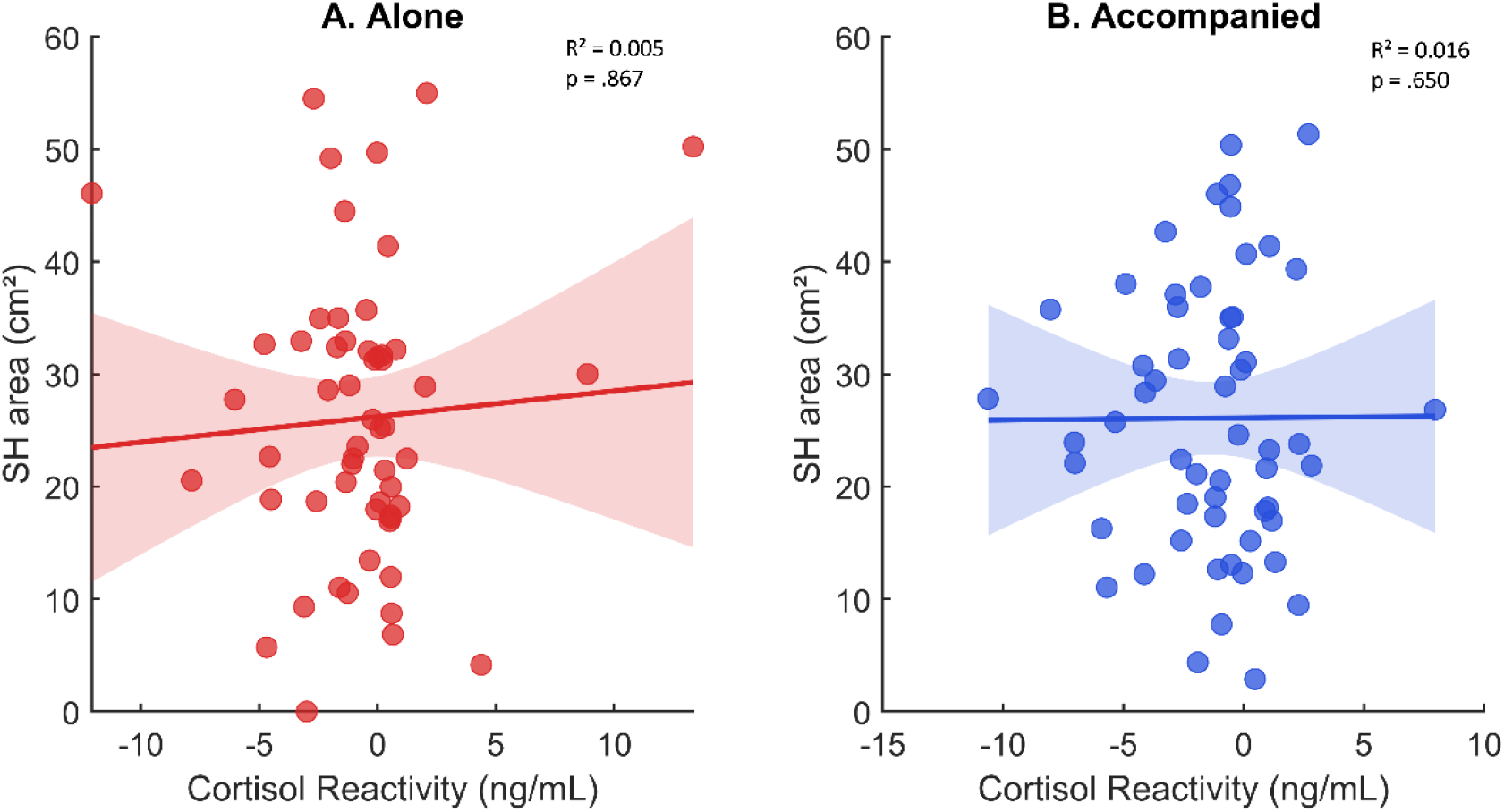
Scatterplots show the association between cortisol reactivity and secondary hyperalgesia area during the alone (A) and accompanied (B) sessions. Each point represents one participant. Regression lines are adjusted for relationship duration, with shaded areas indicating 95% confidence intervals. Red points represent the alone session and blue points represent the accompanied session. R² and *p*-values shown in the figure correspond to the overall multiple regression models including cortisol reactivity and relationship duration as predictors of secondary hyperalgesia area.

#### 3.4.2 Accompanied session

A multiple linear regression was performed to examine whether cortisol reactivity and relationship duration predicted secondary hyperalgesia area during the accompanied session (see figure 4B). The overall model was not significant, *F*(2, 53) = 0.435, *p* = .650, explaining 1.6% of the variance in secondary hyperalgesia area (*R*² = .016, adjusted *R*² = −.021), BF₁₀ =0.15. Cortisol reactivity was not a significant predictor of secondary hyperalgesia area (*B* = −0.017, *SE* = 0.526, β = −0.005, *t*(53) = −0.032, *p* = .974, BF₁₀ =0.27). Relationship duration was also not a significant predictor (*B* = 0.030, *SE* = 0.032, β = 0.127, *t*(53) = 0.933, *p* = .355, BF₁₀ =.39).

### 3.5 Association between partner-related changes in cortisol reactivity and secondary hyperalgesia

A linear regression was conducted to determine whether between-session differences in cortisol reactivity were associated with between-session differences in secondary hyperalgesia, using the difference in secondary hyperalgesia area (alone minus accompanied) as the dependent variable and the between-session difference in cortisol reactivity (alone minus accompanied) as the independent variable, while controlling for relationship duration (see figure 5). The overall model was not significant, *F*(2, 53) = 1.187, *p* = .313, explaining 4.3% of the variance in the between-session difference in secondary hyperalgesia area (*R*² = .043, adjusted *R*² = .007), BF₁₀ =0.27. The between-session difference in cortisol reactivity was not significantly associated with the difference in secondary hyperalgesia area (*B* = −0.595, *SE* = 0.639, β = −0.126, *t*(53) = −0.931, *p* = .356, BF₁₀ =0.43), and relationship duration was also not significantly associated with the outcome (*B* = −0.036, *SE* = 0.033, β = −0.151, *t*(53) = −1.116, *p* = .269, BF₁₀ =0.5).

**Figure 5.**
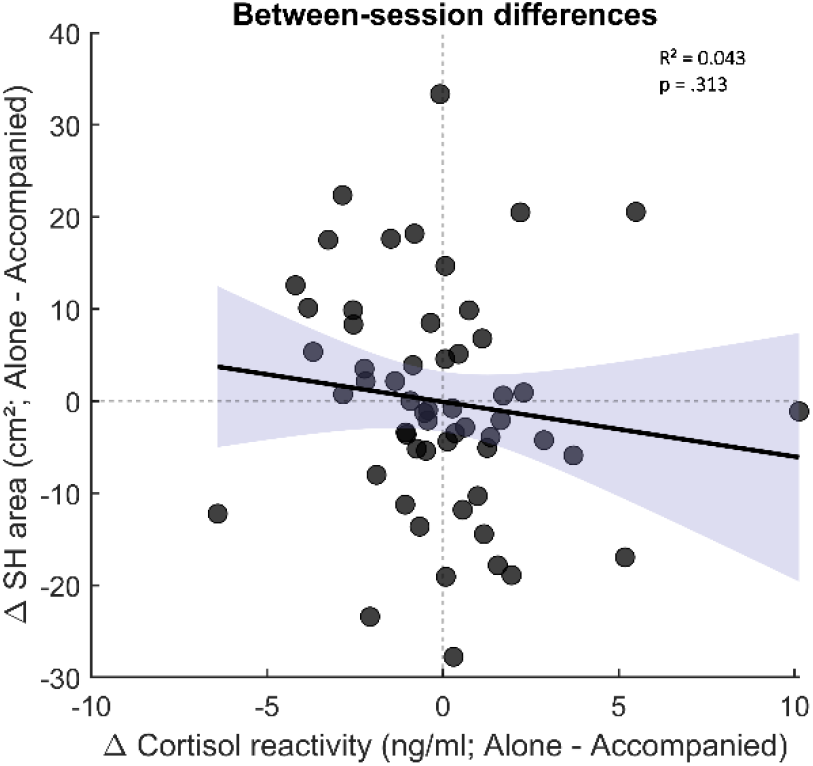
Scatterplot showing the association between the between-session difference in cortisol reactivity and the between-session difference in secondary hyperalgesia area, calculated as alone minus accompanied. Each point represents one participant. The regression line is adjusted for relationship duration, with the shaded area indicating the 95% confidence interval. Dashed reference lines indicate zero difference between sessions. R² and *p*-values shown in the figure correspond to the overall multiple regression model including the between-session difference in cortisol reactivity and relationship duration as predictors of the between-session difference in secondary hyperalgesia area.

## 4. Discussion

The present study investigated whether the presence and touch of a romantic partner modulates salivary cortisol responses during an experimental pain protocol and whether baseline cortisol levels and cortisol reactivity are associated with experimentally induced secondary hyperalgesia. Overall, baseline cortisol levels did not differ between accompanied and alone sessions, and cortisol concentrations decreased from baseline to post-task, with no significant session × sampling time interaction Furthermore, neither baseline cortisol nor cortisol reactivity predicted secondary hyperalgesia, and between-session differences in cortisol reactivity were not associated with differences in secondary hyperalgesia. Together, these findings suggest that salivary cortisol measured before and after the protocol was not associated with secondary hyperalgesia or significantly influenced by partner presence or touch.

Although high-frequency stimulation constitutes a salient nociceptive stimulus, in the present study cortisol concentrations decreased from baseline to the end of the experimental session in both conditions. This finding should be interpreted considering the temporal dynamics of salivary cortisol and the characteristics of our experimental protocol. Moreover, anticipation of an expected stressor has been shown to activate the HPA axis before actual stressor exposure, meaning that cortisol levels obtained before the pain protocol may already reflect participants’ anticipatory responses to the upcoming experimental procedures rather than a true resting baseline (Beltzer, 2022; Engert et al., 2013). In addition, cortisol concentrations typically decline across the waking day following the cortisol awakening response, such that a diurnal decline may also have contributed to the reduction observed between the two sampling points (Dickman et al., 2020). However, given the relatively limited sampling window and the absence of additional cortisol measurements throughout the day, the contribution of the diurnal rhythm cannot be determined in the present study. Because cortisol was sampled only before and after the protocol, the observed changes likely reflect the integrated neuroendocrine response to the entire session, including anticipatory processes, pain induction, affective touch, and progressive adaptation to the experimental context, rather than an isolated response to high-frequency stimulation or the affective touch. This interpretation is further supported by a recent review showing that salivary cortisol is more consistently associated with stress-related processes than with pain intensity itself in experimental settings (Zarnegar et al., 2024). Thus, cortisol reactivity in the present study should be interpreted as reflecting the integrated neuroendocrine response to the experimental context rather than as a direct biomarker of nociceptive processing, consistent with evidence that salivary cortisol is strongly influenced by HPA-axis dynamics, sampling timing, and experimental paradigm (Hellhammer et al., 2009).

Contrary to our hypothesis, changes in cortisol concentrations from baseline to post-task did not differ between the alone and accompanied sessions. This finding contrasts with previous studies reporting attenuated cortisol responses in the presence of social support or during touch by a romantic partner (Berretz et al., 2022; Dreisoerner et al., 2021, 2021; Heinrichs et al., 2003; Hostinar & Gunnar, 2015). However, these studies primarily employed psychosocial stress paradigms, and the neuroendocrine mechanisms engaged during psychosocial stress may differ from those involved during experimentally induced pain. Evidence specifically examining the effects of CT-targeted affective touch on salivary cortisol is more limited. Triscoli et al., (2017) for example, compared CT-optimal touch with vibration and observed no specific effect of CT stimulation on salivary cortisol, with cortisol levels decreasing similarly over time in both groups. Together, these findings suggest that the effects of social touch on HPA-axis activity may depend on the experimental context in which touch is delivered and may not generalize across different experimental paradigms. Furthermore, because cortisol was sampled only before and after the experimental session, transient changes in HPA-axis activity occurring during the protocol may not have been captured.

Neither baseline cortisol levels nor cortisol reactivity predicted the magnitude of secondary hyperalgesia in either the accompanied or the alone session. In addition, the absence of differences in baseline cortisol levels between sessions indicates that participants started both the alone and accompanied sessions with comparable HPA-axis activity, suggesting that the social context did not influence anticipatory cortisol levels prior to pain induction. Although alterations in basal cortisol have been linked to stress-related HPA-axis dysregulation in chronic pain conditions (Vachon-Presseau et al., 2013), such associations were not observed in the present sample of healthy participants. This lack of association is consistent with previous studies reporting heterogeneous relationships between salivary cortisol and experimental pain, likely reflecting differences in stimulus characteristics, sampling protocols, and experimental context (Goodin et al., 2012; Zarnegar et al., 2024, 2025). Furthermore, our previous study using the same experimental protocol and participants sample found that partneŕs presence did not significantly alter the extent of secondary hyperalgesia. The present findings extend those observations by showing that neither baseline cortisol levels nor cortisol reactivity explained individual variability in secondary hyperalgesia, regardless of whether participants completed the protocol alone or in the presence and touch of their romantic partner.

The absence of an association between cortisol and secondary hyperalgesia in the present study contrasts with experimental evidence indicating that glucocorticoid activity can influence pain sensitivity and sensitization. Pharmacological reduction of cortisol has been shown to increase basal mechanical pain sensitivity and temporal summation, whereas acute hydrocortisone administration attenuated secondary hyperalgesia induced by capsaicin (Kuehl et al., 2010; Michaux et al., 2012). However, these effects do not appear to generalize across pain modalities. Acute increases in cortisol were associated with greater sensitivity to visceral pain but did not alter somatic heat pain sensitivity (Benson et al., 2019), while psychosocial stress has been shown to increase pain sensitivity in healthy individuals (Crettaz et al., 2013). More recently, using high-frequency electrical stimulation to induce secondary hypersensitivity, van den Broeke et al., (2026) found that acute stress did not alter the development of hypersensitivity when administered before stimulation, whereas stress applied after hypersensitivity had developed increased pinprick hypersensitivity exclusively in women. Together, these findings suggest that the influence of HPA-axis activity on pain processing may depend not only on the pain modality and sensitization model, but also on the timing of stress or glucocorticoid activation relative to pain induction and sensitization. In contrast to these experimental manipulations, the present study assessed endogenous cortisol without experimentally altering HPA-axis activity, using only pre- and post-task measurements. Thus, the absence of an association between cortisol and secondary hyperalgesia does not necessarily indicate that HPA-axis activity is unrelated to pain sensitization, but rather that cortisol change measured across the entire experimental protocol may lack the temporal resolution to isolate the period in which sensitization develops.

Regarding sex, no significant main effect of sex was observed on cortisol concentrations. However, a significant session × sex interaction indicated that cortisol concentrations differed between the alone and accompanied sessions depending on sex, with women showing higher concentrations in the accompanied session. This interaction was not accompanied by a significant session × sampling time × sex interaction, suggesting that the overall change in cortisol from baseline to post-task did not differ between social contexts as a function of sex. Although these findings provide limited evidence for sex differences in cortisol changes over time, they suggest that sex may contribute to variability in cortisol levels across social contexts. This possibility is consistent with evidence that sex can influence neuroendocrine and pain responses to experimental manipulation. For example, Benson et al., (2019) reported sex-dependent effects of acute hydrocortisone administration on visceral pain sensitivity, whereas somatic heat pain sensitivity was not affected. Men typically show larger cortisol responses to acute stress (Gu et al., 2022; Liu et al., 2017; Zarnegar et al., 2025), but partner support appears to lower cortisol in men and raise it in women (Kirschbaum et al., 1995) although the reverse has also been reported (Berretz et al., 2022). Further studies with larger samples specifically powered to investigate sex differences may help clarify whether sex moderates the relationship between HPA-axis activity, social context, and pain sensitization.

A limitation of the present study is that cortisol was sampled only at two time points. As a result, it was not possible to characterize the temporal dynamics of the HPA-axis response or to distinguish anticipatory activation from pain-related and recovery-related cortisol changes. Additionally, the lack of control over the cortisol awakening response and the natural circadian rhythm of cortisol throughout the day represents a limitation that should be addressed in future investigations. Future studies should include more frequent endocrine sampling to determine whether specific phases of the cortisol response are associated with the development or maintenance of sensitization. Furthermore, to better understand the active components underlying neuroendocrine and pain responses, experimental groups including partner presence without touch, as well as neutral or instrumental touch, would be needed.

## Conclusion

In this study, we investigated whether a romantic partner’s presence and touch modulate salivary cortisol during an experimental pain protocol. Cortisol levels did not differ between the alone and accompanied sessions. Neither baseline cortisol nor cortisol change predicted the extent of secondary hyperalgesia. Overall, our results suggest that salivary cortisol sampled before and after a pain protocol is not sensitive to the presence and touch of a romantic partner and does not capture the mechanisms underlying central sensitization.

## Acknowledgments

This study was conducted at the Psychology Research Centre (CIPsi), School of Psychology, University of Minho, supported by the Foundation of Science and Technology (FCT; UID/01662: Centro de Investigação em Psicologia) through national founds. Márcia da-Silva was supported by a doctoral research grant from the Foundation for Science and Technology (FCT) through the Portuguese State Budget (Ref.:UI/BD/154477/2022). Anabela Silva-Fernandes was supported by the FCT grant (2024.09688.CEECIND). Alberto J. González-Villar was supported by grant RYC2023-044827-I, funded by MICIU/AEI/10.13039/501100011033 and by the FSE+. This work was supported by the EFIC-Grünenthal Grant (E-G-G) 2022.

## Data availability statement

The conditions of our ethical approval do not include the public archiving of anonymized participant data. The cortisol, secondary hyperalgesia, and behavioral data used in this study are available from the corresponding author upon reasonable request, provided that the ethical requirements governing data reuse are maintained.

## REFERENCES

Ackerley, R., Backlund Wasling, H., Liljencrantz, J., Olausson, H., Johnson, R. D., & Wessberg, J. (2014). Human C-Tactile Afferents Are Tuned to the Temperature of a Skin-Stroking Caress. The Journal of Neuroscience, 34(8), 2879–2883. 10.1523/JNEUROSCI.2847-13.2014

Beltzer, E. (2022). An ecological approach for investigations of the anticipatory cortisol stress response. Biological Psychology, 175, 108428. 10.1016/j.biopsycho.2022.108428

Benson, S., Siebert, C., Koenen, L., Engler, H., Kleine-Borgmann, J., Ulrike, B., Icenhour, A., & Elsenbruch, S. (2019). Cortisol affects pain sensitivity and pain-related emotional learning in experimental visceral but not somatic pain: A randomized controlled study in healthy men and women. 160(8), 1719–1728. 10.1097/j.pain.0000000000001579

Berretz, G., Cebula, C., Wortelmann, B. M., Papadopoulou, P., Wolf, O. T., Ocklenburg, S., & Packheiser, J. (2022). Romantic partner embraces reduce cortisol release after acute stress induction in women but not in men. PLOS ONE, 17(5), e0266887. 10.1371/journal.pone.0266887

Bruehl, S., Morris, M. C., & al’Absi, M. (2022). Stress-induced analgesia: An evaluation of effects on temporal summation of pain and the role of endogenous opioid mechanisms. Pain Reports, 7(2), e987. 10.1097/PR9.0000000000000987

Crettaz, B., Marziniak, M., Willeke, P., Young, P., Hellhammer, D., Stumpf, A., & Burgmer, M. (2013). Stress-induced allodynia—Evidence of increased pain sensitivity in healthy humans and patients with chronic pain after experimentally induced psychosocial stress. PloS One, 8(8), e69460. 10.1371/journal.pone.0069460

da-Silva, M., Ribeiro-Carreira, A., Oliveira, M., Sampaio, A., Coutinho, J., & González-Villar, A. J. (2026a). Affective CT-touch attenuates cortical responses and subjective attention during temporal summation of second pain. The Journal of Physiology. 10.1113/JP290421

da-Silva, M., Ribeiro-Carreira, A., Oliveira, M., Sampaio, A., Coutinho, J., & González-Villar, A. J. (2026b). The activation of CT fibers or affective touch does not affect pain sensitization by secondary hyperalgesia (p. 2026.01.27.701964). bioRxiv. 10.64898/2026.01.27.701964

Dickman, K. D., Thomas, M. C., Anderson, B., Manuck, S. B., & Kamarck, T. W. (2020). Social Integration and Diurnal Cortisol Decline: The Role of Psychosocial and Behavioral Pathways. Psychosomatic Medicine, 82(6), 568–576. 10.1097/PSY.0000000000000825

Dreisoerner, A., Junker, N. M., Schlotz, W., Heimrich, J., Bloemeke, S., Ditzen, B., & van Dick, R. (2021). Self-soothing touch and being hugged reduce cortisol responses to stress: A randomized controlled trial on stress, physical touch, and social identity. Comprehensive Psychoneuroendocrinology, 8, 100091. 10.1016/j.cpnec.2021.100091

Engert, V., Efanov, S. I., Duchesne, A., Vogel, S., Corbo, V., & Pruessner, J. C. (2013). Differentiating anticipatory from reactive cortisol responses to psychosocial stress. Psychoneuroendocrinology, 38(8), 1328–1337. 10.1016/j.psyneuen.2012.11.018

Goodin, B. R., Quinn, N. B., King, C. D., Page, G. G., Haythornthwaite, J. A., Edwards, R. R., Stapleton, L., & McGuire, L. (2012). Salivary cortisol and soluble tumor necrosis factor-α receptor II responses to multiple experimental modalities of acute pain. Psychophysiology, 49(1), 118–127. 10.1111/j.1469-8986.2011.01280.x

Gu, H., Ma, X., Zhao, J., & Liu, C. (2022). A meta-analysis of salivary cortisol responses in the Trier Social Stress Test to evaluate the effects of speech topics, sex, and sample size. Comprehensive Psychoneuroendocrinology, 10, 100125. 10.1016/j.cpnec.2022.100125

Heinrichs, M., Baumgartner, T., Kirschbaum, C., & Ehlert, U. (2003). Social support and oxytocin interact to suppress cortisol and subjective responses to psychosocial stress. Biological Psychiatry, 54(12), 1389–1398. 10.1016/s0006-3223(03)00465-7

Hellhammer, D. H., Wüst, S., & Kudielka, B. M. (2009). Salivary cortisol as a biomarker in stress research. Psychoneuroendocrinology, 34(2), 163–171. 10.1016/j.psyneuen.2008.10.026

Hostinar, C. E., & Gunnar, M. R. (2015). Social Support Can Buffer against Stress and Shape Brain Activity. AJOB neuroscience, 6(3), 34–42. 10.1080/21507740.2015.1047054

Jaltare, K. P., Manresa, J. B., Niwa, S., & Torta, D. M. (2024). Verbal Support From a Stranger Reduces the Development of Mechanical Hypersensitivity: Behavioral and Neurophysiological Evidence. The Journal of Pain, 25(10), 104599. 10.1016/j.jpain.2024.104599

Jaltare, K. P., Vanderijst, L., Karos, K., & Torta, D. M. (2023). The impact of the social context on the development of secondary hyperalgesia: An experimental study. PAIN, 164(12), 2711. 10.1097/j.pain.0000000000002971

Kirschbaum, C., Klauer, T., Filipp, S. H., & Hellhammer, D. H. (1995). Sex-specific effects of social support on cortisol and subjective responses to acute psychological stress. Psychosomatic Medicine, 57(1), 23–31. 10.1097/00006842-199501000-00004

Krahé, C., Paloyelis, Y., Condon, H., Jenkinson, P. M., Williams, S. C. R., & Fotopoulou, A. (2015). Attachment style moderates partner presence effects on pain: A laser-evoked potentials study. Social Cognitive and Affective Neuroscience, 10(8), 1030– 1037. 10.1093/scan/nsu156

Kuehl, L. K., Michaux, G. P., Richter, S., Schächinger, H., & Anton, F. (2010). Increased basal mechanical pain sensitivity but decreased perceptual wind-up in a human model of relative hypocortisolism. Pain, 149(3), 539–546. 10.1016/j.pain.2010.03.026

Leone, C. M., Lenoir, C., & van den Broeke, E. N. (2025). Assessing signs of central sensitization: A critical review of physiological measures in experimentally induced secondary hyperalgesia. European Journal of Pain (London, England), 29(3), e4733. 10.1002/ejp.4733

Liu, J. J. W., Ein, N., Peck, K., Huang, V., Pruessner, J. C., & Vickers, K. (2017). Sex differences in salivary cortisol reactivity to the Trier Social Stress Test (TSST): A meta-analysis. Psychoneuroendocrinology, 82, 26–37. 10.1016/j.psyneuen.2017.04.007

Löken, L. S., Wessberg, J., Morrison, I., McGlone, F., & Olausson, H. (2009). Coding of pleasant touch by unmyelinated afferents in humans. Nature Neuroscience, 12(5), 547–548. 10.1038/nn.2312

Marshall, A. G., & McGlone, F. P. (2020). Affective Touch: The Enigmatic Spinal Pathway of the C-Tactile Afferent. Neuroscience Insights, 15, 2633105520925072. 10.1177/2633105520925072

McGlone, F., Wessberg, J., & Olausson, H. (2014). Discriminative and Affective Touch: Sensing and Feeling. Neuron, 82(4), 737–755. 10.1016/j.neuron.2014.05.001

Michaux, G. P. N., Magerl, W., Anton, F., & Treede, R.-D. (2012). Experimental characterization of the effects of acute stresslike doses of hydrocortisone in human neurogenic hyperalgesia models. Pain, 153(2), 420–428. 10.1016/j.pain.2011.10.043

Nikolaeva, E. I., Dydenkova, E. A., Mayorova, L. A., & Portnova, G. V. (2024). The impact of daily affective touch on cortisol levels in institutionalized & fostered children. Physiology & Behavior, 277, 114479. 10.1016/j.physbeh.2024.114479

Quartana, P. J., Buenaver, L. F., Edwards, R. R., Klick, B., Haythornthwaite, J. A., & Smith, M. T. (2010). Pain Catastrophizing and Salivary Cortisol Responses to Laboratory Pain Testing in Temporomandibular Disorder and Healthy Participants. The journal of pain : official journal of the American Pain Society, 11(2), 186. 10.1016/j.jpain.2009.07.008

Triscoli, C., Croy, I., Steudte-Schmiedgen, S., Olausson, H., & Sailer, U. (2017). Heart rate variability is enhanced by long-lasting pleasant touch at CT-optimized velocity. Biological Psychology, 128, 71–81. 10.1016/j.biopsycho.2017.07.007

Vachon-Presseau, E., Roy, M., Martel, M.-O., Caron, E., Marin, M.-F., Chen, J., Albouy, G., Plante, I., Sullivan, M. J., Lupien, S. J., & Rainville, P. (2013). The stress model of chronic pain: Evidence from basal cortisol and hippocampal structure and function in humans. Brain: A Journal of Neurology, 136(Pt 3), 815–827. 10.1093/brain/aws371

van den Broeke, E. N., Deckers, L., De Meutter, Z., O’Neill, F., Verhaegen, N., Kusch, S., Magerl, W., Torta, D. M., & Van Diest, I. (2026). The effect of acute stress on high-frequency electrical stimulation-induced pain and pinprick hypersensitivity in healthy women. Pain, 167(2), 308–319. 10.1097/j.pain.0000000000003790

Van Puyvelde, M., Staring, L., Schaffers, J., Rivas-Smits, C., Groenendijk, L., Smeyers, L., Collette, L., Schoofs, A., Van den Bossche, N., & McGlone, F. (2021). Why do we hunger for touch? The impact of daily gentle touch stimulation on maternal-infant physiological and behavioral regulation and resilience. Infant Mental Health Journal, 42(6), 823–838. 10.1002/imhj.21949

von Mohr, M., Krahé, C., Beck, B., & Fotopoulou, A. (2018). The social buffering of pain by affective touch: A laser-evoked potential study in romantic couples. Social Cognitive and Affective Neuroscience, 13(11), 1121–1130. 10.1093/scan/nsy085

Wingenfeld, K., Wolf, S., Kunz, M., Krieg, J.-C., & Lautenbacher, S. (2015). No effects of hydrocortisone and dexamethasone on pain sensitivity in healthy individuals. European Journal of Pain, 19(6), 834–841. 10.1002/ejp.610

Woolf, C. J. (2011). Central sensitization: Implications for the diagnosis and treatment of pain. Pain, 152(3 Suppl), S2-15. 10.1016/j.pain.2010.09.030

You, D. S., Creech, S. K., & Meagher, M. W. (2016). Enhanced Area of Secondary Hyperalgesia in Women with Multiple Stressful Life Events: A Pilot Study. Pain Medicine, 17(10), 1859–1864. 10.1093/pm/pnw049

Zarnegar, R., Vounta, A., Amrapala, A., & Ghoreishizadeh, S. S. (2025). A study of salivary cortisol and glutamate after the cold pressor task in healthy adults. PeerJ, 13, e19625. 10.7717/peerj.19625

Zarnegar, R., Vounta, A., Li, Q., & Ghoreishizadeh, S. S. (2024). Nociception related biomolecules in the adult human saliva: A scoping review with additional quantitative focus on cortisol. Molecular Pain, 20, 17448069241237121. 10.1177/17448069241237121

